# Perceiving troubles from others: a behavioral and fMRI study

**DOI:** 10.64898/2026.02.22.707243

**Authors:** Sandra Chanraud, Thomas Michelet, Alexandre Zenon, Arnaud Boutin, Arnaud Badets

## Abstract

From a behavioral and neuronal perspective, observational and physical practice conditions have been theorized to be equivalent during motor task learning. However, some paradigms can challenge such a functional equivalence hypothesis. The perception of difficulties experienced by others may play a role in observational learning by allowing learners to partially distance themselves from these episodes, thereby limiting their impact on learning. In contrast, during physical practice, performance difficulties are directly experienced, which may constrain such distancing mechanisms. Indeed, an observer watching a model that uses a wrong physical strategy can ignore erroneous trials in order to preserve action encoding. The main goal of the present study was to prevent such observer avoidance and to test the cognitive and neuronal functional equivalence between the physical and observational practice groups. During this experiment, both groups learned two motor sequences. Only one sequence was repeatedly interrupted to perturb encoding. Behavioral results revealed that both groups were equally negatively impacted by such interruptions. Together, these findings suggest that while physical and observational practice can lead to comparable behavioral outcomes under strong disruption, they rely on partially distinct neural strategies. Physical practice predominantly engages motor and striato-cerebellar feedback loops, whereas observational learning relies more strongly on fronto-cerebellar and episodic memory networks, highlighting a context-dependent functional equivalence between learning modalities.

## 1 – Introduction

Understanding difficulties exhibited by others is a core component of several cognitive functions, including empathy, learning, and behavioral adaptation (Bulgarelli & Jones, 2023). For example, in the domain of adaptation, it has been crucial for the different countries of Europe to understand clearly the conflict between Ukraine and Russia. This understanding has enabled different countries to pursue improved defense and rearmament to avoid future conflict with a hypothetical belligerent. In the domain of empathy, Cao and colleagues (2024) suggested that the cognitive mechanism called “empathic pain”, defined as the understanding of an observer about another person’s pain, can be a key component for care behaviors and avoidance responses for future potential disagreements. Finally, in the domain of learning, it could be argued that when a person is confronted with an unknown skill to perform, such a requirement can afford an unpleasant task difficulty to attain the mandatory level of this new skill. In the same vein, observers can also understand the same level of difficulty when they are prompted to learn this skill only by observation (Badets & Blandin, 2010; Han et al., 2022). The present study investigates whether the cognitive and neural mechanisms supporting motor sequence learning are similar or distinct between observational and physical practice when learning is challenged by encoding difficulties.

At a neurophysiological level, numerous neuroimaging experiments investigating the human action-observation system reported that a set of common neural structures is activated during both motor execution or physical practice and action observation, thus providing evidence of a relative “functional equivalence” between practice modalities (Grèzes & Decety, 2001; Mattar & Gribble, 2005; Hardwick et al., 2018). Importantly, these findings suggest that functional equivalence between observation and execution is relative rather than absolute, and may depend on task demands, learning context, and the nature of the cognitive constraints imposed during acquisition. Evidence indicates that observation consistently recruits a set of frontoparietal and posterior temporal regions, within the so-called “action observation network” (AON; Apsvalka, Cross & Ramsey, 2018; Cross et al., 2009; Abreu et al., 2012). The AON encompasses a distributed cortical network of brain regions involved in visual analysis of action, as well as areas involved in visuomotor and sequence learning (Cross et al., 2009; Hardwick et al., 2018), including mainly premotor-parietal and occipital regions but without a consistent recruitment of subcortical areas. More specifically, a large meta-analysis by Hardwick et al. (2018) revealed an extensive overlap between brain regions active during action observation and motor execution, involving a distributed cortical network spanning premotor, parietal, and sensorimotor regions. However, the authors also highlighted differences in the brain networks active during observation and practice. The cortical regions more consistently associated with observation than practice are mainly recruited in the premotor, parietal and occipital cortex. In contrast, practice has been shown to recruit more sensorimotor cortical areas than observation, within a distributed brain network including mainly premotor, motor and somatosensory cortical regions (Hardwick et al., 2018) as well as subcortical structures such as the cerebellum, thalamus and basal ganglia (Haggard, 2008; Henschke & Pakan, 2023).

This theoretical conception of a functional equivalence between action execution and observation at the neurophysiological level is also consistent with behavioral experiments. At a behavioral level, the search for similarities and differences in the cognitive mechanisms engaged during observational and physical practice has been well documented (see Henschke & Pakan, 2023; Rudisch et al., 2024 for recent reviews). For example, in exploring these similarities/differences between the two practice conditions, it has been shown that physical practice involved many more motor components like limbs and their associated muscular coordination (Shea et al., 2000). In contrast, for observational practice, such conditions engaged preferentially an abstract representation of the action to be performed (Badets & Blandin, 2010; see also Schütz-Bosbach & Prinz, 2007 for a similar theoretical account). In order to test this hypothesis, Badets and Blandin (2010) contrasted two different levels of knowledge of results (KR; which is information on the performed outcome) for learners. The first one, called “bandwidth KR”, consisted of giving a quantitative KR on the performed action when participants surpass a predefined bandwidth around the goal of the task. When performance is inside the predefined bandwidth, such a KR is not delivered. Clearly, participants understood this “no-KR” trial as a good performance (i.e., a qualitative KR; see Lee & Carnahan, 1990 for the seminal discovery). To resume, this KR schedule was considered a very powerful tool for learning because it provides both quantitative and qualitative feedback. The second schedule is called “KR frequency”, and it consists of giving KR only for some trials during the acquisition phase, like for 50% of trials, for example. This procedure affords only one quantitative feedback for some trials and necessitates participants to manage their behavior and process their intrinsic feedback during the no-KR trials. Results revealed that the bandwidth KR benefited equally for physical and observational practice conditions. In comparison to a control procedure without bandwidth, the groups under bandwidth KR performed better on a motor sequence task during delayed tests. However, only observers were positively impacted by the KR frequency. For authors, this more “subtle” KR schedule allowed the abstract representation of the task to be much more strongly encoded because of the no-KR trials. Indeed, it is well known that a KR frequency improves the encoding of perceptual features (Winstein & Schmidt, 1990; Blandin, Toussaint & Shea, 2008), leading to an enhanced abstract representation of the action. The trials without KR are assumed to improve the perceptual process that controls the overall coordination of the motor task, and to a greater extent during observational practice when no motor execution is required.

From another but close theoretical point of view, such involvement of an abstract representation throughout observational practice is also in line with the associative sequence-learning hypothesis, which suggests that the link between sensory and motor features is likewise flexible (Cook, Bird, Catmur, Press, & Heyes, 2014; Heyes, 2001). Badets and colleagues (2018) supported this conclusion using a “Stop” paradigm for motor sequence learning through observational and physical practice. The main goal of this experiment was to introduce a “trouble” at encoding to test whether observers and practitioners could manage this difficulty through similar strategies. Using a dyad-like protocol, participants were randomly assigned to a physical or observational practice group. All participants were required to learn two motor sequences (sequences A and B) either by action observation or physical practice. During the acquisition phase, the participant “observer” had to learn the two motor sequences only by observing the “practitioner”. However, during this learning phase, sequence B was interrupted by the message “STOP” displayed on the screen to instruct the practitioner to stop practicing the motor sequence. This stop signal afforded a strong disappointment for participants in both groups, and it was expected that the learning of motor sequence B would be hindered compared to the learning of motor sequence A. The results showed that only the practitioners were negatively affected by the stop signal, leading to impaired learning of the interrupted sequence B compared to sequence A. In the observational practice group, however, both the uninterrupted sequence A and the interrupted sequence B were equally learned. This suggests that, during the learning phase, observers may have adopted a safety strategy allowing them to develop and preserve an efficient abstract representation of the interrupted motor sequence without suffering from this interruption of the model’s performance. According to the authors, and in line with the flexible hypothesis of the associative sequence-learning theory (Cook et al., 2014; Heyes, 2001), observational learning enabled participants to build an abstract representation of the task, which in turn permitted an equivalent encoding of both motor sequences.

However, it could be argued that the stop paradigm used by Badets and colleagues (2018) constituted only a “subtle” experimental manipulation, which may have facilitated the emergence of an abstract representation for observers. Indeed, in this stop paradigm, the participants were interrupted just once. Specifically, only one stop signal occurred during a practice block in which participants performed the sequence. It can therefore be hypothesized that repeated stop signals would induce a stronger negative effect in observers, which in turn would affect the encoding of sequence B similarly to participants in the physical practice group. Hence, an extended “stop” paradigm would likely make the functional equivalence between physical and observational practice more apparent.

In the present experiment, the stop signal was embedded within the sequence, producing a stronger interruption effect within a block of trials. From this perspective, we hypothesized that a similar “stop” effect would emerge in both the observation (OBS) and physical practice (PP) groups, as behaviorally expressed by better performance on sequence A than sequence B. At the neurophysiological level, our primary goal was to determine whether this expected similar level of behavioral performance for the OBS and PP groups is underpinned by the engagement of a common or specific, partially overlapping neural network. However, even if OBS and PP share a relative functional equivalence at the cognitive and neural levels, research devoted to identifying the specificity of observational learning has revealed that OBS may rely on distinct neurocognitive processes than those triggered by PP (Hardwick et al., 2018), sometimes leading to different behavioral outcomes (Badets & Blandin, 2010; Badets, Boutin, & Michelet, 2018; Conessa et al., 2023; Hayes, et al., 2012). Hence, it can also be hypothesized that OBS and PP would behave differently. According to the safety hypothesis for observational learning, observers may adopt a safety strategy that allows them to learn both motor sequences effectively, irrespective of the amount of exposure to the interrupted sequence. This would imply that interruptions do not interfere with motor sequence learning, and thus are reflected by the involvement of distinct neural networks and cognitive processes between practice modalities.

## 2 – Methods

### 2.1 – Participants

We estimated the sample size based on the previous behavioral study on the “stop-practice” paradigm during physical and observational learning (Badets et al., 2018; *N* = 14 per group). Accordingly, we recruited 34 students (*N* = 17 per group) from the University of Bordeaux (*mean age*: 21.2 years; *SD* = 2 years; 19 females). None of them had prior knowledge/skill with the motor sequence. Each participant was invited to read and sign an informed consent form about the task and the experimental procedure. This study was realized in agreement with the ethical principles of the 1964 Helsinki Declaration under the number CPP 18-INSB-01.

### 2.2 – Apparatus, task, and General procedure

We used the apparatus and task developed by Badets and colleagues (2018). Specifically, participants completed the experiment in front of a computer screen, either inside the scanner (during several phases) or in the experimental room (for one phase). On each trial, four horizontally aligned white empty squares were presented in the center of the screen. These squares corresponded to the spatial locations of the response keys (1, 2, 3, and 4) on the response box placed on the participant’s abdomen inside the scanner, or on a standard keyboard positioned on a table in the experimental room. Each imperative stimulus consisted of one of the four squares filled in black. Participants were requested to press with their four right-hand fingers (without the thumb) the appropriate response key as rapidly and accurately as achievable in order to perform and encode two second-order conditional (SOC) 12-element motor sequences (keys for Sequences A: 4-1-3-2-1-4-2-3-4-3-1-2, and Sequence B: 2-4-2-1-3-4-1-2-3-1-4-3). Each trial began with the presentation of a fixation cross for 1000 ms, followed by the display of four empty squares for a random interval between 2000 and 4000 ms. Immediately after this presentation, the first imperative stimulus appeared. The participant’s response triggered the presentation of the next stimulus until the achievement of the entire sequence.

For the general procedure, participants were tested using a dyadic protocol (see Table 1). They were requested to perform three main phases: two MRI sessions (pre-test & post-test) and an intermediate learning phase conducted outside the MRI scanner in the experimental room. The recording of responses was managed by the E-prime software (version 3.0; Psychology Software Tools, Inc.) for the MRI sessions and the MATLAB R2017b software (MathWorks, Natick, NA) for the learning phase conducted outside the MRI scanner. Specifically, participants attended first an MRI session which included anatomical and resting-state acquisitions, followed by a BOLD acquisition during the pre-learning phase (pretest), during which the participant had to start learning sequences A and B. Imaging data were acquired using a 3T Siemens Prisma MRI (Siemens Healthcare, Germany) equipped with a 64-channel head/neck coil. Participants were placed in a supine position in the MRI scanner, and their heads were immobilized with cushions to reduce motion artifacts. The visual stimuli were projected onto a translucent screen located behind the scanner bore, and participants looked at the screen via a mirror fixed inside the scanner at approximately 20 cm above their eyes. The response keypad was located comfortably on the participant’s belly.

**Table 1.** Experimental groups and procedures. The three experimental phases (pre-test, acquisition, post-test) are reported for each experimental group (PP and OBS group). PP: Physical practice; OBS: Observational practice.

| Experimental groups and procedures |  |  |  |
| --- | --- | --- | --- |
|  | Pre-test<br>(30 minutes) | Acquisition<br>(40 minutes) | Post-test<br>(30 minutes) |
| **Groups** |  |  |  |
| **PP** | Anatomical<br>and resting-state acquisitions<br>*5 minutes rest*<br>+ MRI on the sequences A<br>(20 trials) & B (20 trials) | PP for the sequences A<br>(100 trials) & B (100<br>trials) | Anatomical<br>and resting-state acquisitions<br>*5 minutes rest*<br>+ MRI on the sequences A<br>(20 trials), B (20 trials), and<br>C (20 trials) |
| **OBS** | Identical Procedure | OBS for the sequences A<br>(100 trials) & B (100<br>trials) | Identical Procedure |

Anatomical images were obtained with a 3D MPRAGE T1-weighted MRI sequence with the following parameters: FA = 9°; Inversion Time = 1070 ms, repetition time = 2000 ms, echo time = 2 ms, field of view (FOV) = 256 × 256 mm2 to cover the entire brain, 192 slices, and an isotropic voxel size = 1 × 1 × 1 mm3. Six hundred resting-state dynamic scans were acquired using a 2D simultaneous multi-slice echo gradient echo planar sequence with the following parameters: 2.5 × 2.5 mm voxels in-plane; 2.5 mm slice thickness with no gap; 60 transverse slices, FOV = 210 × 210 mm; Matrix = 84 x 84; TR = 700 ms; TE = 30 ms; multiband slice acceleration factor of 6; phase encoding direction Anterior-Posterior; flip angle 53°; bandwidth 2705 hz/pixel; effective echo spacing 0.49 ms. During the functional MRI session, participants were instructed to remain eyes-closed and not to fall asleep. A whole-brain coverage, multiband EPI acquisition was used for the learning phases in the scanner: TR= 700 ms; TE = 30 ms; FOV = 210×210 mm2; 60 slices, 360 dynamics for each run, voxel size = 2.5 mm3. All acquisitions were realigned along the anterior commissure–posterior commissure (AC-PC) plane.

Inside the scanner, both participants of the dyad received the same instructions to perform the task. While one participant was completing the pre-test (and the same for the post-test), their practice-group counterpart waited outside the scanner room, and vice versa (order counterbalanced across participants). The first MRI phase served to familiarize participants with the task. The pre– and post-tests each consisted of performing 40 trials, with 20 trials on Sequence A and 20 trials on Sequence B. Each trial consisted of the presentation of one 12-element motor sequence (i.e., 12 successive stimuli). The presentation of the sequence A or B was alternated and counterbalanced across the participants. For the post-test, after the completion of the sequence A and B, the participants completed 20 trials on a new motor Sequence C (2-4-1-3-2-1-4-2-3-4-3-1). This transfer test served as an index of sequence-specific learning. Indeed, if performance is equivalent between this new sequence C and the Sequences A and B, then no evidence of sequence-specific learning for Sequences A and B could be inferred (see Krakauer et al., 2019, for a review).

During the second phase (acquisition), participant in the OBS group stood upright and behind the PP participant, located above the right shoulder to apprehend the visual information from the right-hand finger movements and sequential stimuli on the screen. The dyad was requested to learn the motor Sequences A and B (10 trials per block; in total 10 blocks per sequence), and was also informed that for some trials, Sequence B would be interrupted by a stop signal. The stop signals appeared inside a trial in a faded schedule across blocks: Blocks 1-2 contained no stop trials; Blocks 3-4 contained 2 stop trials (occurring after the fourth press); Blocks 5-6 contained 4 stop trials; Blocks 7-8 contained 6 stop trials; and Blocks 9-10 contained 8 stop trials. Within each block, stop trials were presented in a random order across participants.

Between phases, we counterbalanced the performance of Sequences A and B across dyads. Specifically, some dyads began the pre– and post-tests with Sequence A, while others began with Sequence B. Similarly, some participants began the acquisition phase with Sequence A, and others with Sequence B. Sequence order was also counterbalanced between the pre-/post-test phases and the acquisition phase.

After the last scanning session, participants filled out a recall questionnaire on both interrupted and uninterrupted sequences (see Boutin et al., 2014; Destrebecqz & Cleeremans, 2001; Destrebecqz et al., 2005). This recall test assessed explicit sequence knowledge. Specifically, participants were requested to write down the sequential order of the 12 elements that composed the learned motor Sequences A and B on a sheet of paper from memory.

### 2.4 – Dependent variables for motor performance

At the behavioral level, the dependent variables in this study were: (1) the response time, which represents the time between the imperative stimuli and the response keys; (2) the error rate, which represents the number of erroneous key presses; and (3) the paper-and-pencil score, which represents the number of sequence elements correctly recalled.

### 2.5 – fMRI data processing

The resting-state functional images were preprocessed using SPM12 (https://www.fil.ion.ucl.ac.uk/spm/software/spm12/) and the CONN toolbox (https://web.conn-toolbox.org/). For each participant, the first five scans were discarded, then the functional images were realigned to the middle scan to correct for small head movements that might have occurred during the acquisition. A mean functional image was created at this step, onto which the anatomical image was coregistered and segmented, enabling the later use of an anatomical gray matter image for subsequent normalization of functional images. Then, the gray matter image was warped to the MNI template (MNI 152, Montreal Neurological Institute, McGill University, Montreal, Canada) in order to define the normalization parameters, which were applied to the functional images. The normalized functional images were then smoothed with a 5-mm full width at half maximum (FWHM) Gaussian kernel. The functional images were then detrended to correct for the loss of signal along the acquisition, and signals from the white matter and cerebrospinal fluid regions were extracted using principal component analysis. Functional volumes were then band-pass filtered (0.009-0.08 Hz) to restrict analyses within the frequency band of interest. Finally, the functional images were despiked to remove outliers from the time-series signal.

The functional BOLD activation images acquired during the pre– and post-tests were preprocessed as described above, except that detrending and despiking were not applied. Also, in the activation images, signals from white matter and cerebrospinal fluid were not regressed out; instead, a gray matter mask was used for the analyses.

### 2.6 – FMRI test activation sequence analyses

Preprocessed data were analyzed using a general linear model (GLM; Friston et al., 1994). Each sequence was defined as a separate regressor, resulting in 4 regressors (preA; postA; preB; postB). Contrasts were estimated for each effect of interest (i.e., postA vs preA; postB vs preB). To estimate the changes between conditions, we used a paired-samples t-test design. Then for each group separately, a random-effects analysis was performed using a one-sample t-test for each contrast image to obtain population-level inferences for each condition in both groups. All results were thresholded at p<0.001 (uncorrected) and a cluster size > 10 contiguous voxels.

### 2.7 – Resting state statistical analyses

All connectivity analyses were run using the CONN toolbox (version 19, MIT, Cambridge, MA, USA). The mean BOLD signal was first extracted from a parcellated brain of 116 regions (Automated Anatomical Labeling (AAL) atlas (Tzourio-Mazoyer et al., 2002). Then, a functional connectivity (FC) matrix of correlations among the BOLD signals extracted in the AAL regions was constructed for the resting-state (Rs) acquisitions. Whole-brain changes in Rs FC values (ΔFC) associated with sequence learning were quantified by subtracting the Rs-FC values of the acquisition done after the post-test (Rs2) from those acquired before the pretest (Rs1), resulting in Δ values (Rs2 – Rs1). These ΔFC values were used as a within-subject variable in the two following analyses run in the CONN toolbox. A contrast between the OBS and the PP groups was carried out, equivalent to an ANOVA, to identify pairs of regions for which the FC differed between the two groups. All results were thresholded at p<0.001 (uncorrected).

## 3 – Results

### 3.1 – Response time analysis

To assess whether participants in both groups correctly encoded the motor Sequence A and were impacted by the stop paradigm, we analyzed response times with a multivariate analysis of variance (ANOVA) with the following factors: 2 *Groups* (PP and OBS) X 2 *Tests* (pre-test, post-test) X 2 *Sequences* (A and B) with repeated measures for the last two factors. Groups served as the between-participants factor, and Tests and Sequences as the within-participants factors. The analysis revealed a main effect of the factor *Tests*, *F*(1, 32) = 101.4, *p* < .0001, *Sequences*, *F*(1, 32) = 28.8, *p* < .0001, and most importantly a *Tests* X *Sequences* interaction*, F*(1, 32) = 4.9, *p* < .03, *η*^2^ = .13. The *Groups* X *Tests* X *Sequences* interaction was not significant, (*F*(1, 32) < 1). For the *Tests* X *Sequences* interaction, Scheffé post-hoc analysis revealed that participants of both groups did not differ in performance on the motor sequences A and B during the pre-test, but exhibited reduced performance on the motor sequence B in comparison to the sequence A during the post-test (pre-test: Sequence A (564±96 ms) and Sequence B (576±104 ms) / post-test: Sequence A (439±101 ms) and Sequence B (467±101 ms)). Noticeably, both groups have improved their performances from the pre-test to the post-test for both motor sequences. For the sake of clarity, the Table 2 presents the mean response times for each group, test, and sequence.

**Table 2.** Behavioral results – Response time. Mean response times (ms) and standard deviations (ms; in brackets) are specified for each experimental group. Letter A, B, and C specifies the motor sequence A, B, and C, respectively.

| Groups | Experimental groups and response time data |  |  |
| --- | --- | --- | --- |
|  | Pre-test | Post-test | Transfer test |
| Action observation | A: 550 (90)<br>B: 567 (100) | A: 444 (95)<br>B: 470 (89) | C: 498 (85) |
| Physical practice | A: 578 (103)<br>B: 586 (110) | A: 433 (109)<br>B: 464 (114) | C: 500 (105) |

To determine whether the sequences A and B have been specifically learned in comparison to the sequence C, we analyzed response times of the post-test using an ANOVA with the following factors: 2 G*roups* (PP and OBS) X 3 *Sequences* (Sequences A, B, and C) with repeated measures on the last factor. Groups served as the between-participants factor, and Sequences as the within-participants factor. The analysis revealed only a main effect of the factor *Sequences*, *F*(2, 64) = 27.6, *p* < .0001, *η*^2^ = .46. Scheffé post-hoc analysis revealed that participants of both groups performed better on Sequence A than on Sequences B and C, and better on Sequence B than on Sequence C (Sequence A: 439±101 ms; Sequence B: 467±101 ms; Sequence C: 499±94 ms). This finding confirms the stop effect, as evidenced by poorer performance on Sequence B relative to Sequence A. More importantly, it also demonstrates sequence-specific learning, as both practiced sequences (A and B) were performed better than the unpracticed Sequence C, indicating that learning was specific to the trained sequences rather than reflecting a general improvement in task performance. Finally, the factor *Groups* and *Groups X Sequences* interaction were not significant, *F*(1, 32) < 1; *F*(2, 64) < 1, respectively.

To assess whether participants of the PP group have improved their performances on the Sequences A and B during the acquisition phase, we analyzed response times using a repeated-measures ANOVA with the following factorial design: *10 Blocks* (10 blocks of 10 trials each) X 2 S*equences* (A and B). Blocks and Sequences served as within-participants factors. The analysis revealed a main effect of the factor *Sequences*, *F*(1, 16) = 6.35, *p* < .02, *η2* = .28, indicating that participants performed better on Sequence A (363±42) than on Sequence B (386±32 ms). α main effect of the factor *Blocks* was also observed, *F*(9, 144) = 18.27, *p* < .0001, *η2* = .53, reflecting significant performance improvements across practice blocks. Scheffé post-hoc comparisons revealed that participants improved significatively their performances from the first block (Block 1: 421±61 ms) to the last block (Block 10: 323±84 ms) during the acquisition phase. To illustrate this learning curve, and for the sake of clarity, we present only an example from the block 1 (421±61) to the block 10 (323±84). The *Blocks* X *Sequences* interaction was not significant, *F*(9, 144) = 1.77, *p* < .08. Finally, participants performed the motor sequences during this acquisition phase with very few errors (2,2%).

### 3.2 – Error rate analysis

For the following sections, similar multivariate ANOVAs were performed on the error rates for the pre– and post-test. For the first experimental plan (2 G*roups* (PP and OBS) X 2 *Tests* (pre-test, post-test) X 2 S*equences* (A and B), with repeated measures for the last two factors), the analysis revealed a main effect of the factor *Tests*, *F*(1, 32) = 4.4, *p* < .04, *Sequences*, *F*(1, 32) = 8.2, *p* < .007, and most importantly a *Tests* X *Sequences* interaction*, F*(1, 32) = 6.6, *p* < .01, *η*^2^ = .17. The *Groups* X *Tests* X *Sequences* interaction was not significant, (F(1, 32) < 1). For the *Tests* X *Sequences* interaction, Scheffé post-hoc analysis revealed that participants in both groups performed equally on the motor Sequences A and B during the pre-test (2.12% vs. 2.21%, respectively) but exhibited more errors on Sequence B in comparison to Sequence A during the post-test (3.35% vs. 2.09%, respectively).

For the second experimental plan (2 G*roups* (PP and OBS) X 3 *Sequences* (Sequences A, B, and C), with repeated measures on the last factor), the analysis revealed only a significant main effect of the factor *Sequences*, *F*(2, 64) = 9.5, *p* < .0002, *η*^2^ = .23. Scheffé post-hoc analysis revealed that participants in both groups performed better on Sequence A (2.09%) than on Sequences B (3.35%) and C (3.78%). Importantly, the error rate on Sequence B was not significantly different from that of Sequence C, suggesting that the stop effect negatively impacted performance for both groups. Finally, the factor *Groups* and *Groups X Sequences* interaction were not significant, *F*(1, 32) < 1; *F*(2, 64) = 1.06, *p* < .35, respectively.

### 3.3 – Explicit sequence knowledge

To compare explicit knowledge of the two sequences (A and B), scores from the paper-and-pencil tests were submitted to an ANOVA with the following factors: G*roups* (PP and OBS) X *Sequences* (A and B) with repeated measures for the last factor. The analysis revealed no significant main effects of *Groups*, *F*(1, 32) = 0.16, *p* = .68, or *Sequences*, *F*(1, 32) = 1.59, *p* = .22, and no *Groups* X *Sequences* interaction, *F*(1, 32) = 0.05, *p* = .81. It is important to note that participants of both groups recalled more than five keys for each of the two sequences (PP and OBS: 7 and 6 elements for Sequences A and B, respectively). In this perspective, it can be assumed that the encoding of the two motor sequences relied mostly on an explicit mechanism, since a recall of more than five elements from a SOC 12-element motor sequence is considered an indicator of conscious-explicit sequence knowledge (Badets et al. 2023; Willingham & Goedert-Eschmann, 1999).

### 3.4 – Learning-related activations

One-sample t-tests revealed significant activations for the uninterrupted contrast (Sequence A: post-test vs. pre-test) and the interrupted contrast (Sequence B: post-test vs. pre-test) in both groups (see Figure 1; Tables 3-6).

**Figure 1.**
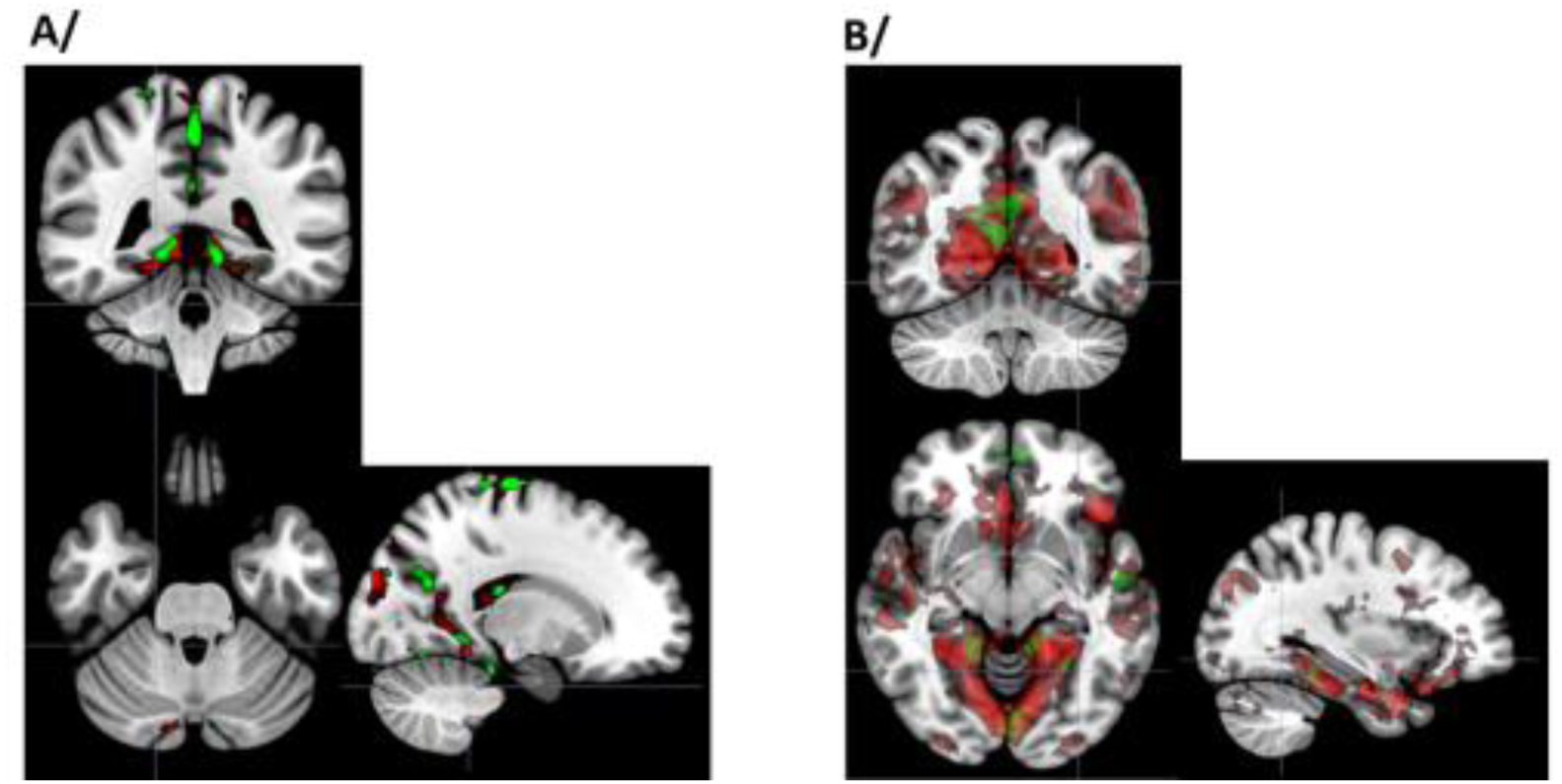
Functional activations in the contrast post-test versus pre-test for. A/ the uninterrupted motor sequence and B/ the interrupted motor sequence. Clusters of activations are represented in red for the Physical Practice group and in green for the Observational group.

**Table 3.**
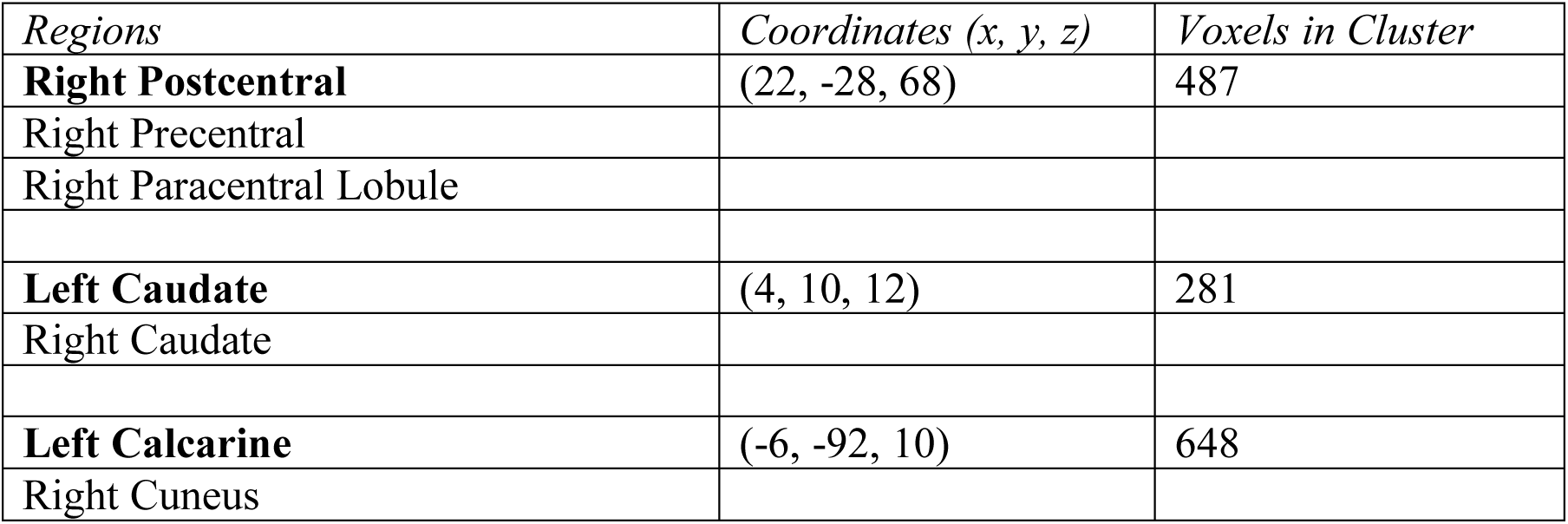
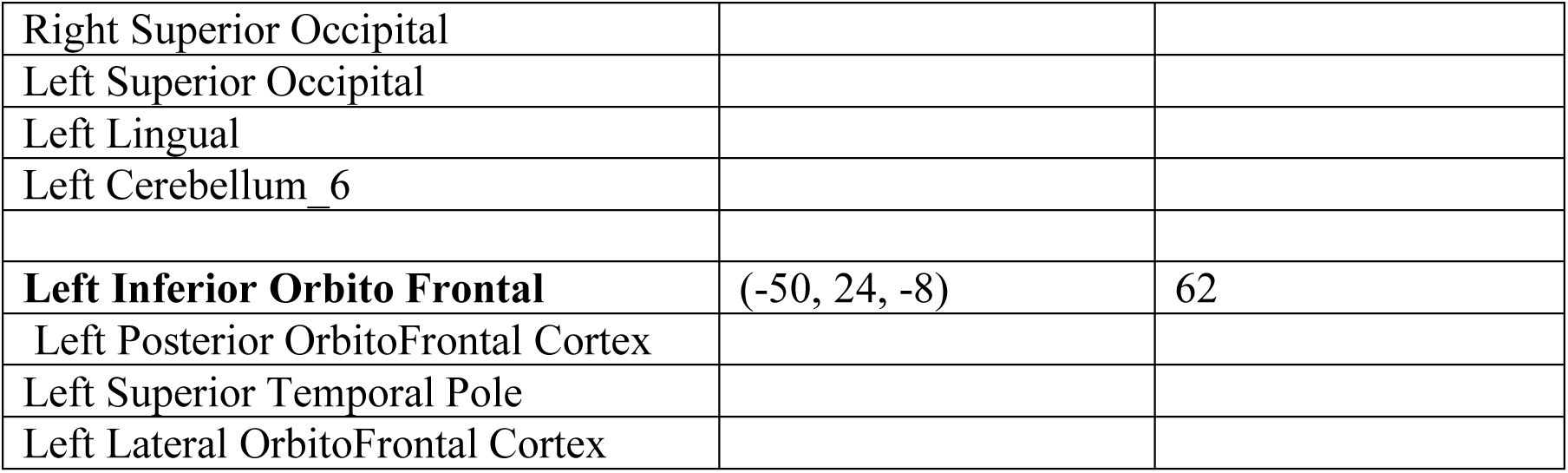
Functional activations in the contrast post-test versus pre-test for the uninterrupted motor sequence. A in the PP group. P <0.001 uncorrected. Minimum cluster size = 10 voxels. x-, y– and z-coordinates are specified in the MNI (Montreal Neurological Institute) space.

Brain Activation Tables
| *Regions* | *Coordinates (x, y, z)* | *Voxels in Cluster* |
| --- | --- | --- |
| **Right Postcentral** | (22, -28, 68) | 487 |
| Right Precentral |  |  |
| Right Paracentral Lobule |  |  |
| **Left Caudate** | (4, 10, 12) | 281 |
| Right Caudate |  |  |
| **Left Calcarine** | (-6, -92, 10) | 648 |
| Right Cuneus |  |  |

|  |  |  |
| --- | --- | --- |
| Right Superior Occipital |  |  |
| Left Superior Occipital |  |  |
| Left Lingual |  |  |
| Left Cerebellum_6 |  |  |
| **Left Inferior Orbito Frontal** | (-50, 24, -8) | 62 |
| Left Posterior OrbitoFrontal Cortex |  |  |
| Left Superior Temporal Pole |  |  |
| Left Lateral OrbitoFrontal Cortex |  |  |

## Brain Activation Tables

### 3.5 – Between-groups learning-induced functional connectivity changes

The between-groups comparison for the Rs2>Rs1 contrast revealed significant FC differences in a set of pairs of regions (Figure 2). The analysis revealed two distinct patterns of FC changes between the groups:

(i). The ΔFC increased more in the PP group in comparison to the OBS group between the left Caudate and the left Hippocampus (T(32) = −4.07), and between the right Caudate and the right Parahippocampus (T(32) = –3.77).
(ii). The ΔFC increased more in the OBS group than in the PP group between the right Supplementary Motor Area and the left Crus I (T(32) = 3.97), between the right Cerebellum IV-V and the left Fusiform (T(32) = 3.65) and between the right middle Frontal cortex and the Vermis VIII (T(32) = 3.64).

**Figure 2.**
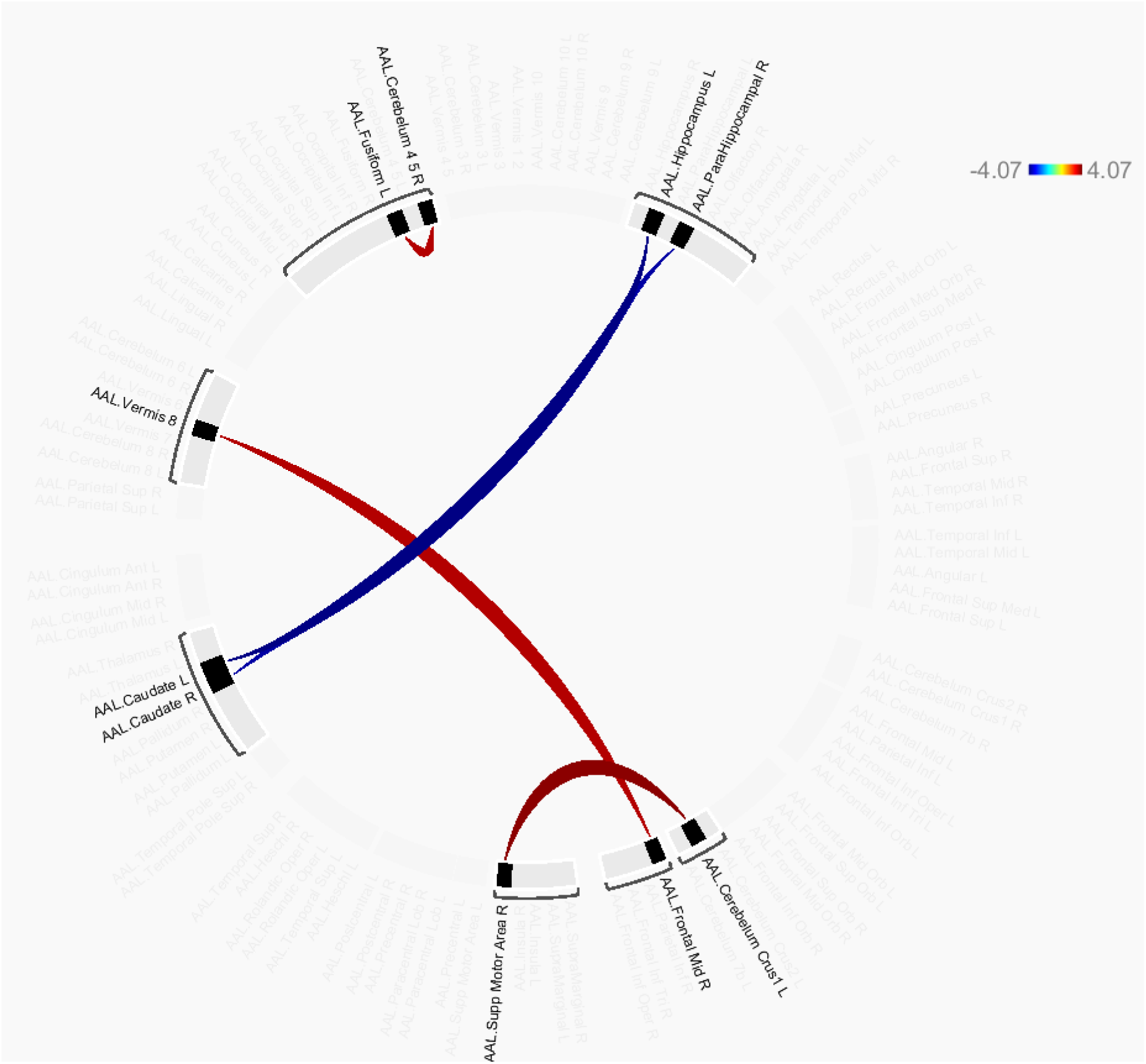
Between-groups differences in functional connectivity changes between the resting session following the learning phases (Rs2) and the resting session before the learning phases (Rs1). Blue connections indicating stronger post-training increases in the Physical Practice group, and red connections indicating stronger post-training increases in the Observational group. Darker color indicates stronger correlation coefficient between brain regions.

## 4 – Discussion

The present study sets out to examine whether the neural mechanisms underlying motor sequence learning differ between physical practice (PP) and observational learning (OBS) when the task involves unexpected disruptions. In line with the “safety hypothesis” of observational learning (Badets et al., 2018), observers could have been less affected by interruptions due to their ability to construct and preserve an abstract representation of the task. However as hypothesized, when more interruptions are introduced during the acquisition phase, both behavioral and neuroimaging results revealed a more nuanced picture.

The present study provides three main contributions. First, it demonstrates that sufficiently frequent interruptions disrupt motor sequence learning in both physical and observational practice. Second, it shows that similar behavioral impairments can arise from distinct neural mechanisms. Third, it refines the concept of functional equivalence by demonstrating its context-dependent nature.

As indicated by the improvements in motor performance from the pre-test to the post-test, participants in both groups have learned the two motor sequences (see the section *Response time analysis*). Importantly, both PP and OBS groups showed reduced performance on the interrupted sequence (Sequence B) compared to the uninterrupted one (Sequence A), as reflected in slower response times and higher error rates. This behavioral outcome suggests that sufficiently frequent stop signals equally disrupted the encoding process in both groups, thereby challenging previous findings that observers are immune to such disturbances (Badets et al., 2018). It is therefore likely that the increased number of interruptions (stop signals) in the present protocol amplified their disruptive effect, narrowing the motor performance gap between the physical and observational practice groups.

The current neuroimaging findings provide compelling evidence for modality– and context-specific neural mechanisms underlying motor learning. The BOLD activation analyses confirm classic predictions for PP. During the execution of Sequence A (uninterrupted), practitioners recruited the pre– and post-central cortices, the paracentral lobule, and cerebellar regions (Table 3). This network reflects strong engagement of motor and sensorimotor circuits underpinning overt movement execution. The involvement of the cerebellum (lobule VI in particular) and primary motor cortex aligns with their well-established roles in timing coordination and procedural consolidation (Doyon et al., 2009; Doyon et al., 2018; Hardwick et al., 2013). In addition, orbitofrontal activation suggests contributions of effort regulation and motivational control during task execution.

For the interrupted sequence, participants in the PP group additionally recruited visual regions (cuneus and lingual gyri), the ventral striatum, cerebellar regions and the precuneus (Table 4). These additional brain activations likely reflect an increased demand for online correction, error monitoring, and effortful adjustments in response to interruptions. The involvement of striatal and visual areas aligns with feedback-driven monitoring models (Ullsperger et al., 2014), in which the motor system intensifies sensory and striatal loops to sustain performance.

**Table 4.**
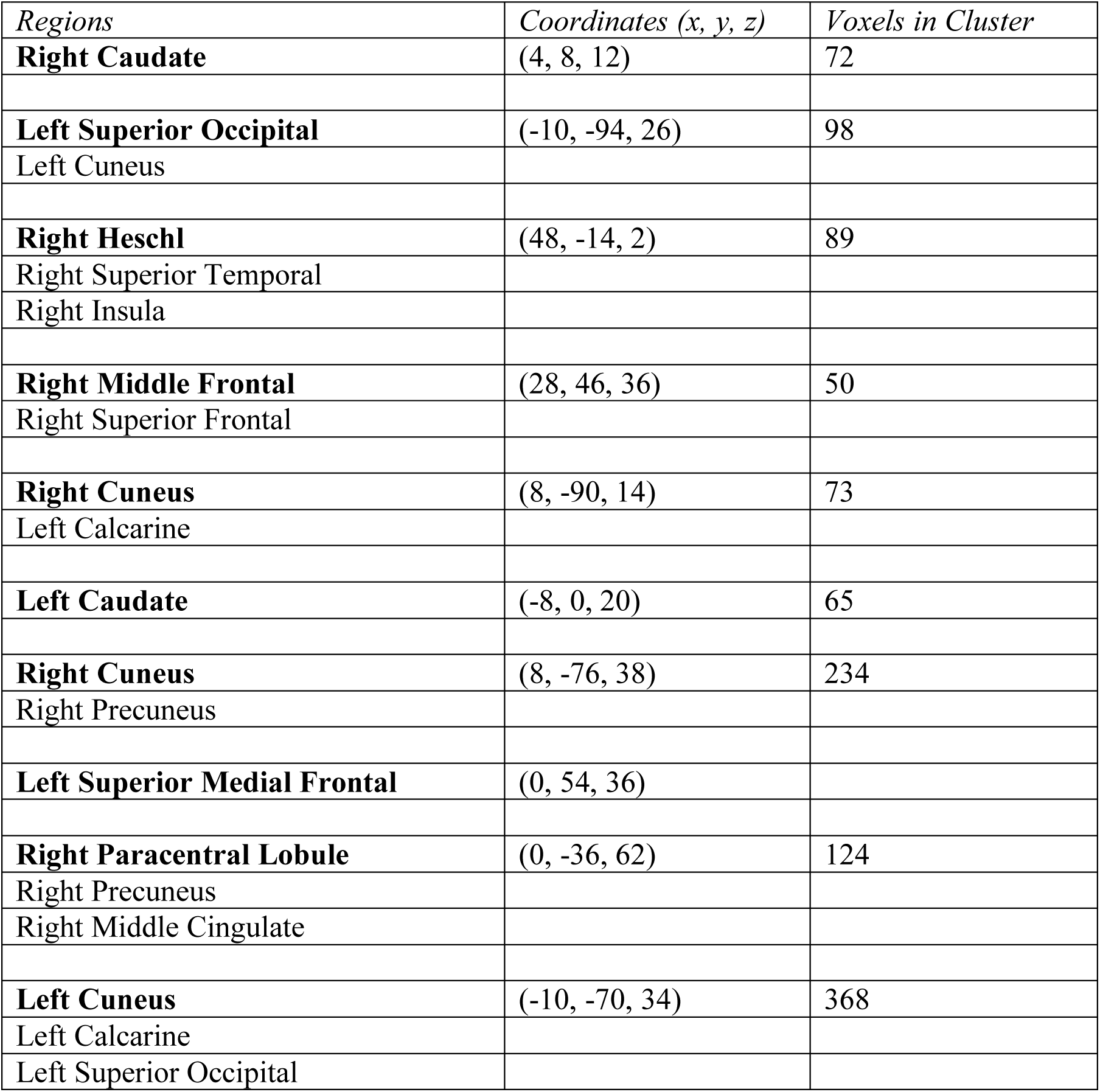
Functional activations in the contrast post-test versus pre-test for the uninterrupted motor sequence. A in the OBS group. P <0.001 uncorrected. Minimum cluster size = 10 voxels. x-, y– and z-coordinates are specified in the MNI space.

| *Regions* | *Coordinates (x, y, z)* | *Voxels in Cluster* |
| --- | --- | --- |
| **Right Caudate** | (4, 8, 12) | 72 |
| **Left Superior Occipital** | (-10, -94, 26) | 98 |
| Left Cuneus |  |  |
| **Right Heschl** | (48, -14, 2) | 89 |
| Right Superior Temporal |  |  |
| Right Insula |  |  |
| **Right Middle Frontal** | (28, 46, 36) | 50 |
| Right Superior Frontal |  |  |
| **Right Cuneus** | (8, -90, 14) | 73 |
| Left Calcarine |  |  |
| **Left Caudate** | (-8, 0, 20) | 65 |
| **Right Cuneus** | (8, -76, 38) | 234 |
| Right Precuneus |  |  |
| **Left Superior Medial Frontal** | (0, 54, 36) |  |
| **Right Paracentral Lobule** | (0, -36, 62) | 124 |
| Right Precuneus |  |  |
| Right Middle Cingulate |  |  |
| **Left Cuneus** | (-10, -70, 34) | 368 |
| Left Calcarine |  |  |
| Left Superior Occipital |  |  |

Resting-state connectivity analyses further support this interpretation. Following learning, practitioners showed increased connectivity between the caudate and hippocampus, as well as between the caudate and parahippocampus. This FC pattern is coherent with procedural consolidation through striato-hippocampal circuits, which integrate sequence memory and motor skill acquisition (Doyon & Benali, 2005). Altogether, the PP condition appears to rely on direct motor integration, feedback-based correction, and consolidation within subcortical, motor-specialized networks.

In contrast, OBS participants exhibited a markedly different brain activation profile. For the uninterrupted Sequence A, brain activations were observed in the cuneus and temporal regions including the insula (Table 5). This network highlights reliance on visual processing and motor simulation via the action observation network (Caspers et al., 2010). In contrast to PP, there was minimal activation of the primary motor cortex, suggesting that observational learning depends more heavily on perceptual processing and internal simulation without overt execution. Additionally, the involvement of the superior frontal cortex may reflect working memory and planning demands to maintain the sequence in an abstract representational form.

**Table 5.** Functional activations in the contrast post-test versus pre-test for the interrupted motor sequence in the PP group. P <0.001 uncorrected. Minimum cluster size = 10 voxels. x-, y– and z-coordinates are specified in the MNI space.

| *Regions* | *Coordinates (x, y, z)* | *Voxels in Cluster* |
| --- | --- | --- |
| **Right Caudate** | (4, 2, 18) | 529 |
| Left Caudate |  |  |
| **Right Precentral** | (12, -28, 78) | 232 |
| Right Postcentral |  |  |
| Right Paracentral Lobule |  |  |
| **Right Calcarine** | (14, -52, 6) | 125 |
| Right Lingual |  |  |
| Vermis 4 5 |  |  |
| Right Cerebellum 4 5 |  |  |
| Right Precuneus |  |  |
| **Right Olfactory** | (2, 20, 0) | 52 |
| Left Olfactory |  |  |
| Left Ventral Striatum |  |  |
| **Left Precuneus** | (-2, -42, 56) | 99 |
| Right Precuneus |  |  |
| Right Paracentral Lobule |  |  |
| Left Paracentral Lobule |  |  |
| **Right Cuneus** | (-6, -84, 38) | 189 |
| Calcarine |  |  |
| **Left Cuneus** | (-4, -86, 34) | 63 |
| Left Superior Occipital |  |  |
| **Right Lingual** | (0, -80, -4) | 156 |
| Left Lingual |  |  |
| **Left Lingual** | (-30, -46, -4) | 100 |
| Left Fusiform |  |  |
| Left Cerebellum 4 5 |  |  |
| Left Precuneus |  |  |

**Table 6.** Functional activations in the contrast post-test versus pre-test for the interrupted motor sequence in the OBS group. P <0.001 uncorrected. Minimum cluster size = 10 voxels. x-, y– and z-coordinates are specified in the MNI space.

| *Regions* | *Coordinates (x, y, z)* | *Voxels in Cluster* |
| --- | --- | --- |
| **Right Middle Temporal** | (52, -4, -26) | 357 |
| Right Middle Pole Temporal |  |  |
| Right Superior Temporal |  |  |
| Right Inferior Temporal |  |  |
| Right Superior Pole Temporal |  |  |
| Right Parahippocampal |  |  |
| **Left Caudate** | (-6, -6, 20) | 674 |
| Right Caudate |  |  |
| Right Middle Cingulate |  |  |
| Left Middle Cingulate |  |  |
| Right Hippocampus |  |  |
| Right Caudate |  |  |
| **Left Lingual** | (-10, -44, -8) | 87 |
| Left Cerebellum Crus 1 |  |  |
| Left Fusiform |  |  |
| Left Parahippocampal |  |  |
| **Left Middle Temporal** | (-52, -10, -8) | 73 |
| Right Middle Temporal |  |  |
| **Right Fusiform** | (36, -26, -18) | 144 |
| Right Parahippocampal |  |  |
| Right Cerebellum 4 5 |  |  |
| Right Hippocampus |  |  |
| **Right Precuneus** | (20, -64, 32) | 319 |
| Left Cuneus |  |  |
| Right Cuneus |  |  |
| Left Precuneus |  |  |
| Left Superior Occipital |  |  |
| Right Superior Occipital |  |  |
| Left Precentral |  |  |
| Left Lingual |  |  |
| **Left Fusiform** | (-28, -48, -12) | 200 |

For the interrupted Sequence B in the OBS group, brain activations extended to the middle and inferior temporal regions, parahippocampus, hippocampus, and precuneus (Table 6). This profile indicates a different strategy: rather than amplifying motor correction processes, observers appeared to rely on episodic and memory-related mechanisms. The joint involvement of the hippocampus and precuneus points to episodic encoding of the disrupted sequence, perhaps as a series of perceptual events rather than as a continuous motor plan. Such a profile aligns with the associative sequence-learning theory (Heyes, 2001; Cook et al., 2014), which emphasizes flexible coupling between perceptual cues and motor representations.

Resting-state functional connectivity further underscores this distinction. Unlike PP learners, observers showed increased connectivity between the medial frontal cortex, cerebellar Crus I and lobules IV-V, vermis, and visual areas. This fronto-cerebellar configuration is more consistent with associative learning and cognitive control, suggesting that early consolidation following observational learning relies primarily on attention and memory networks rather than striato-cerebellar motor loops. The absence of striatal connectivity strengthens the interpretation that OBS consolidates experiences at a perceptual-memory level rather than through procedural memory circuits.

Together, these findings refine the notion of functional equivalence between execution and observation (Grèzes & Decety, 2001; Mattar & Gribble, 2005). Both modalities support sequence learning and are affected by interruptions, but they rely on distinct neural circuits. PP engages sensorimotor integration and striato-cerebellar feedback loops, while OBS relies on perceptual processing, episodic memory, and fronto-cerebellar networks. This divergence does not invalidate the idea of shared neurofunctional mechanisms but emphasizes that equivalence is relative and modulated by task demands—in this case, the frequency of interruptions.

An intriguing interpretation is that interruptions drive practitioners toward enhanced motor correction, while pushing observers toward episodic encoding strategies. Thus, the same disruptive stimulus induces distinct adaptive strategies depending on practice modality. This perspective enriches our understanding of motor learning by showing that observational learning does not merely passively mimic execution but engages complementary, more abstract and memory-dependent processes.

Several limitations should be acknowledged. First, interpretations related to “motivation” or “abstraction” are inferred from activations in brain regions such as the frontal cortex, hippocampus, or precuneus. While these associations are plausible, they should be confirmed through complementary measures (e.g., motivational questionnaires, episodic memory tests). Second, the absence of longitudinal follow-up assessments prevents conclusions about potential differences in retention over time between PP and OBS. Finally, because the task involved explicit learning processes (as indexed by recall scores), it remains an open question whether more implicit sequence learning would yield the same pattern of results.

In conclusion, this study demonstrates that frequent interruptions impair learning in both PP and OBS groups, challenging the notion that observers are fully immune to disruption. However, the two practice groups acquire the interrupted motor sequence through distinct neural strategies: PP recruits motor and striatal feedback loops for correction, while OBS engages episodic memory and fronto-cerebellar networks. These findings emphasize that observational learning is not a passive substitute for physical practice but relies on complementary cognitive and neural mechanisms. Understanding how task constraints modulate the balance between motor, perceptual, and memory systems may have important implications for motor rehabilitation and skill training protocols. Future work on motor learning should further explore how motor, perceptual, and memory systems interact across practice modalities.

## Acknowledgments

Océane Thierre, Yann Gaussens, Elisa Le Cozannet, Lilou Murroni, Julie Goddard for their help in recruitment, data acquisition, collection and analyses.

## Compliance with Ethical Standards

The authors declare that they have no conflict of interest.

All procedures performed in this study were in accordance with the ethical standards of the institutional and national research committee, and with the 1964 Helsinki declaration. Finally, informed consent was obtained from all individual participants included in the study.

## Data availability

The data that support the results of this study are available from the corresponding author upon reasonable request.

